# SARS-CoV-2-specific T cells in unexposed adults display broad trafficking potential and cross-react with commensal antigens

**DOI:** 10.1101/2021.11.29.470421

**Authors:** Laurent Bartolo, Sumbul Afroz, Yi-Gen Pan, Ruozhang Xu, Lea Williams, Chin-Fang Lin, Elliot S. Friedman, Phyllis A. Gimotty, Gary D. Wu, Laura F. Su

**Affiliations:** Department of Medicine, Division of Rheumatology, Perelman School of Medicine, Institute for Immunology, University of Pennsylvania, Philadelphia, PA 19104, USA; Corporal Michael J Crescenz VA Medical Center, Philadelphia, PA, 19104, USA; Division of Gastroenterology and Hepatology, Perelman School of Medicine, University of Pennsylvania, Philadelphia, PA 19104, USA; Department of Biostatistics, Informatics, and Epidemiology, Perelman School of Medicine, University of Pennsylvania, Philadelphia PA 19104, USA

**Author notes:** indicates co-first authors (Laurent Bartolo, Sumbul Afroz, Yi-Gen Pan, Ruozhang Xu). Corresponding author and lead contact: Laura F. Su, University of Pennsylvania, 421 Curie Blvd BRBII/III 311, Philadelphia, PA 19104.

## Abstract

The baseline composition of T cells directly impacts later response to a pathogen, but the complexity of precursor states remains poorly defined. Here we examined the baseline state of SARS-CoV-2 specific T cells in unexposed individuals. SARS-CoV-2 specific CD4^+^ T cells were identified in pre-pandemic blood samples by class II peptide-MHC tetramer staining and enrichment. Our data revealed a substantial number of SARS-CoV-2 specific T cells that expressed memory phenotype markers, including memory cells with gut homing receptors. T cell clones generated from tetramer-labeled cells cross-reacted with bacterial peptides and responded to stool lysates in a MHC-dependent manner. Integrated phenotypic analyses revealed additional precursor diversity that included T cells with distinct polarized states and trafficking potential to other barrier tissues. Our findings illustrate a complex pre-existing memory pool poised for immunologic challenges and implicate non-infectious stimuli from commensal colonization as a factor that shapes pre-existing immunity.

**One Sentence Summary:** Pre-existing immunity to SARS-CoV-2 contains a complex pool of precursor lymphocytes that include differentiated cells with broad tissue tropism and the potential to cross-react with commensal antigens.

## Introduction

SARS-CoV-2 infection (COVID-19) presents with a myriad of organ involvement, variable duration of symptoms, and diverse clinical presentations that ranged from asymptotic infection to death *(1-6)*. The underlying determinants for variable responses to COVID-19 and other infections remain incompletely understood. Age, sex, co-morbidities, and host genetics have emerged as key factors that increase the risk for severe COVID-19 *(7-12)*. Studies in mice have additionally demonstrated a key role of environment on pathogen resistance, with an accumulation of memory T cells providing protection against infections in mice raised in a free-living environment *(13-15)*. What remains less clear is how the environment shapes human immune responses to SARS-CoV-2 and other pathogens. We and others have previously reported that memory T cells recognize human immunodeficiency virus (HIV), cytomegalovirus (CMV), herpes simplex virus (HSV), and yellow fever virus (YFV) in people who tested negative for these infections *(16-18)*. Prior studies have also identified T cells that recognized SARS-CoV-2 in unexposed individuals *(15, 19-25)*. The association between pre-existing cells and protective features such as milder disease and abortive infection further raised questions about the composition of pre-existing pool and how this type of immunity is generated *(26-28)*. A widely held view is that pre-existing memory to SARS-CoV-2 reflects antigen experiences from past exposures to common circulating coronaviruses (CCCoV) *(29)*. Consistent with this, a portion of T cells that respond to SARS-CoV-2 also responds to similar peptides from CCCoV *(30, 31)*. However, other studies have also identified SARS-CoV-2 reactive T cells that lack cross-reactivity to CCCoV antigens *(32)*. The presence of pre-existing memory T cells that recognize viruses without closely related circulating relatives further suggests the possibility that sources other than similar pathogens could contribute to precursor T cell differentiation.

Here we performed an in-depth analysis of SARS-CoV-2 specific T cells in unexposed individuals and investigated alternative sources of antigens that could drive pre-existing T cell differentiation. Rare SARS-CoV-2 specific CD4^+^ T cells were identified directly ex vivo using class II peptide-MHC (pMHC) tetramer enrichment to minimize phenotypic perturbations. Blood samples collected prior to year 2020 were used to ensure the absence of prior SARS-CoV-2 exposure. Hypothesizing that the imprints of initial antigen engagement are retained in pre-existing T cells, we used trafficking molecules to gain insights into where prior antigen engagement might have occurred. We showed that a subset of pre-existing SARS-CoV-2 specific memory CD4^+^ T cells displayed gut homing markers and cross-reacted with commensal-derived antigens in a MHC-dependent manner. Integrated phenotypic analyses revealed additional diversity in the pre-existing states that included cells with tropism to other barrier tissues, including the skin. A portion of SARS-CoV-2 precursors also displayed markers characteristics of regulatory T cell (Treg), T follicular helper cell (Tfh), and Th1 cells. These data highlight the complexity of SARS-CoV-2 specific T cells at baseline and implicate the contribution of microbiome in the development of pre-existing immunity.

## Results

### Pre-existing SARS-CoV-2 specific T cells in unexposed adults

We analyzed PBMC from twelve unexposed donors using pre-pandemic blood samples collected prior to year 2020 (Table S1). SARS-CoV-2 specific CD4^+^T cells were identified using a direct ex vivo approach with peptide-MHC (pMHC) tetramers. We recombinantly expressed HLA-DRA/DRB1*0401 monomers (DR4). A set of 12 peptides that stimulated T cells in COVID-19 patients based on prior studies and predicted to bind HLA-DR4 were selected for generation of tetramers (Table S2). We coupled tetramer staining with magnetic column-based enrichment to enable enumeration of rare tetramer-labeled T cells in the unprimed repertoire. The baseline differentiation states of tetramer-labeled T cells were delineated by anti-CD45RO and CCR7 antibody staining. In total, we analyzed 117 SARS-CoV-2 specific CD4^+^ populations in 12 healthy unexposed adults. SARS-CoV-2 precursors were detectable in samples from all donors but showed large inter-individual and antigen-dependent variations. (Fig. 1A-B, Fig. S1A). On average, the frequency of spike-specific T cells was lower compared to cells that recognized epitopes outside of the spike region (S1 region: 0.7(+/-1.1), S2 region: 0.7 (+/- 0.6), non-spike region: 4.0 (+/- 3.5) cells/million CD4^+^ T cells, Fig. S1C).

**Figure 1:**
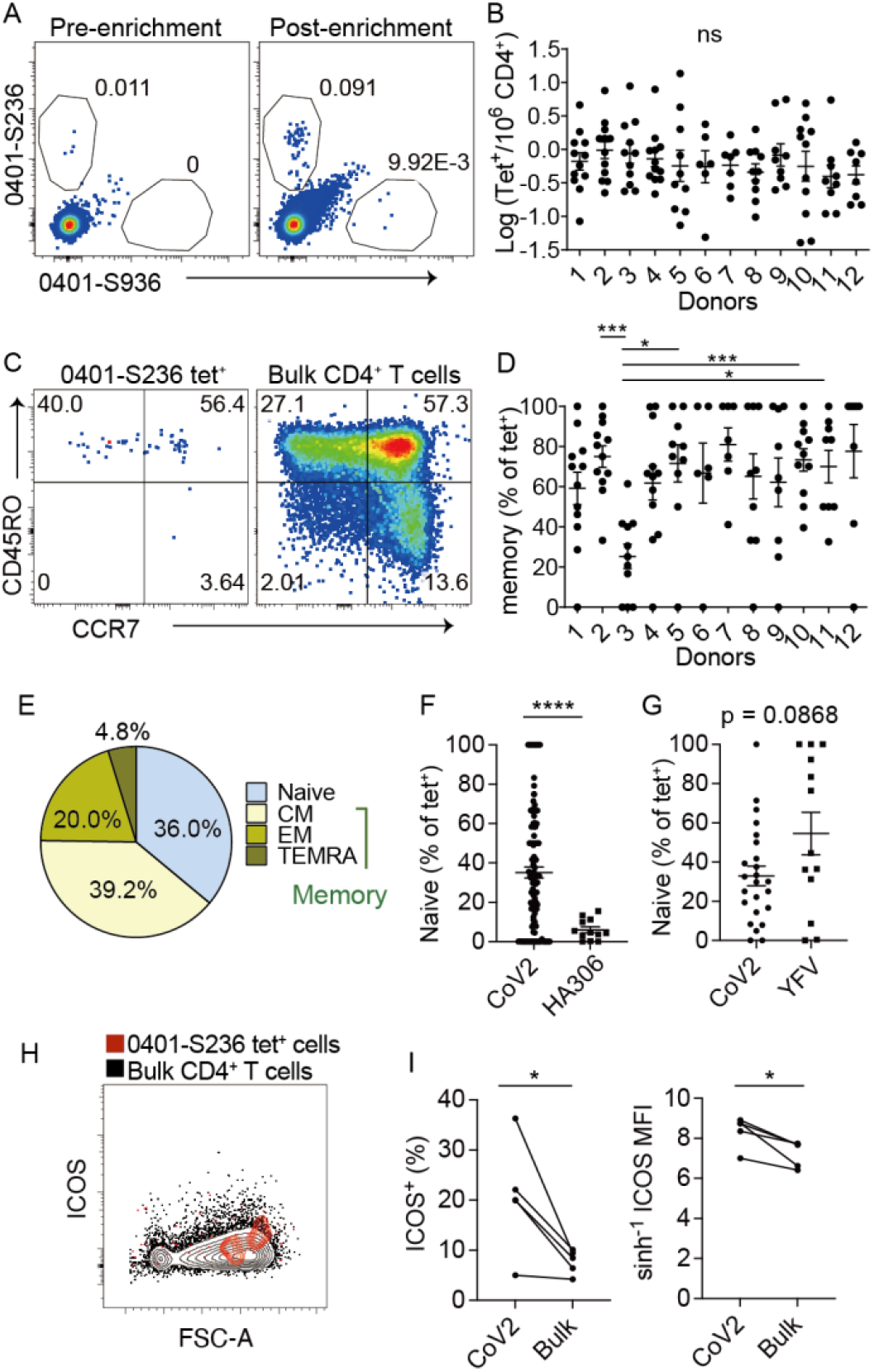
SARS-CoV-2 specific T cells in unexposed donors. (A) Direct ex vivo staining, pre- and post-magnetic enrichment, of representative SARS-CoV-2 tetramer^+^ cells using cryopreserved cells obtained before year 2020. (B) The frequency of SARS-CoV2-specific CD4^+^ T cells across 12 unexposed donors by direct ex vivo tetramer staining. Each symbol represents data from a distinct SARS-CoV-2-specific population, repeated 2.2 times (± 0.2). (C) Plots show anti-CD45RO and CCR7 staining that divide cells into naïve or memory subsets. Bulk CD4^+^ T cells are shown for comparison (right). (D) The abundance of pre-existing memory T cells as a percentage of tetramer^+^ cells shown in B. Pre-existing memory include all cells that do not have a naïve phenotype. (E) Differentiation phenotype of tetramer^+^ cells (n = 117 populations): naïve (CD45RO^-^CCR7^+^), central memory (CM, CD45RO^+^CCR7^+^), effector memory (EM, CD45RO^+^CCR7^-^), and TEMRA (CD45RO^-^CCR7^-^). (F-G) Plot compares the percentage of naïve cells within the indicated specificities. Each symbol represents a tetramer-labeled population. For (F), data combine SARS-CoV-2 and HA306-specicfic T cells from all 12 donors. Data in (G) are limited to two donors who had YFV-specific T cell results from a previous study. (H) Representative plot showing ICOS staining on tetramer^+^ and background CD4^+^ T cells. (I) ICOS expression in SARS-CoV2 tetramer^+^ cells or bulk CD4^+^ T cells as a percentage (left) or by median staining intensity (MFI, right). Distinct SARS-CoV-2 tetramer^+^ populations from the same donor are combined and represented as an average. Each symbol represents data from one individual (n = 5). Line connects data from the same donor. (B) and (D) used Welch’s ANOVA. P-values for pairwise comparisons were computed using Dunnett’s T3 procedure. For (F) and (G), Welch’s t-test was performed. For (I), a paired t-test was performed. Data are shown as Mean ± SEM. * p < 0.05, *** p < 0.001. **** P < 0.0001.

Next, we broadly divided tetramer-labeled cells by CD45RO and CCR7 staining to examine precursor differentiation states. This revealed heterogeneous memory phenotypes that differed by epitope specificity and varied across donors (Fig. 1C-D, S2A). Collectively, only 36% of cells expressed a naïve phenotype. The remaining cells expressed a memory phenotype, which included 39.2% central memory cells (CM), 20% effector memory cells (EM), and 4.8% terminal effector cells (TEMRA) (Fig. 1E). Although naïve SARS-CoV-2 specific T cells comprised of a minor subset, they were still substantially more abundant than the naïve component of hemagglutinin antigen (HA)-specific T cells that have experienced antigens from past exposures to vaccines and/or infections (Fig. 1F). To examine how SARS-CoV-2 specific T cells differ from a true antigen-inexperienced population, we compared them to T cells that recognized YFV in YFV naïve individuals *(18)*. Unlike coronaviruses, YFV-related viruses are uncommon in the United States. This comparison showed less naïve cells in a portion of SARS-CoV-2 specific populations (Fig. 1G). While this finding likely reflects prior exposures to other coronaviruses, we did not find a clear relationship between the abundance of SARS-CoV-2 memory precursors and the conservation of epitopes with CCCoV sequences (Fig. S2B, Table S3). Instead, the abundance of pre-existing memory cells correlated with donor age, suggesting a contribution from the broader antigen experiences that accumulate over time (Fig.S2C). To investigate if T cells are receiving stimulatory signals at baseline, we analyzed activation state using ICOS *(18, 33)*. We found ICOS expressed on a subset of SARS-CoV-2 tetramer^+^ cells (Fig. 1H). The overall ICOS signal was also higher on SARS-CoV-2 specific T cells compared to background CD4^+^ T cells within the same individuals (Fig. 1I). Thus, pre-immune SARS-CoV-2 specific T cells have variable frequencies and display a range of differentiation phenotypes in healthy adults. Our data suggest that acquisition of pre-existing memory states could involve antigens beyond conserved epitopes from other coronaviruses.

### Pre-existing SARS-CoV-2 specific T cells express gut homing markers

The intestinal tract is home to trillions of microbial organisms *(34)*. The composition of the microbiome is heavily dependent on the living environment and has critical impact on human health *(13, 35)*. Additionally, previous studies have demonstrated that homeostatic interactions with microbes in the gut are sufficient to drive human memory T cell differentiation *(36)*. We therefore hypothesized that the gut-microbiome could be a source of constant antigenic stimulation that shapes the pre-existing T cell repertoire. Flexible engagement of TCRs may then allow some commensal-activated T cells to be detected as memory cells to a new pathogen. To begin to test this idea, we used gut homing receptors, integrin β7 and CCR9, to infer priming by and/or the potential to engage intestinal antigens *(37-39)*. Tetramer staining for SARS-CoV2-specific T cells was combined with antibody staining for trafficking receptors. HA-specific T cells were identified for comparison. This revealed gut-tropic SARS-CoV-2 specific T cells that expressed integrin β7 and CCR9 in a coordinated pattern (Fig. 2A-B). Integrin β7 and CCR9 expression were highly variable, with levels that differed between T cell specificities within and across individuals (Fig. 2C-D). While no single SARS-CoV-2 specificity emerged to consistently express gut trafficking receptors, significantly more SARS-CoV-2-specific cells expressed an integrin β7^+^CCR9^+^ phenotype compared to influenza-specific T cells or bulk memory CD4^+^ T cells from the same donor (Fig. 2E). Taken together, these data support the possibility of antigen engagement in the intestinal environment for a subset of SARS-CoV-2 specific pre-existing memory T cells.

**Figure 2:**
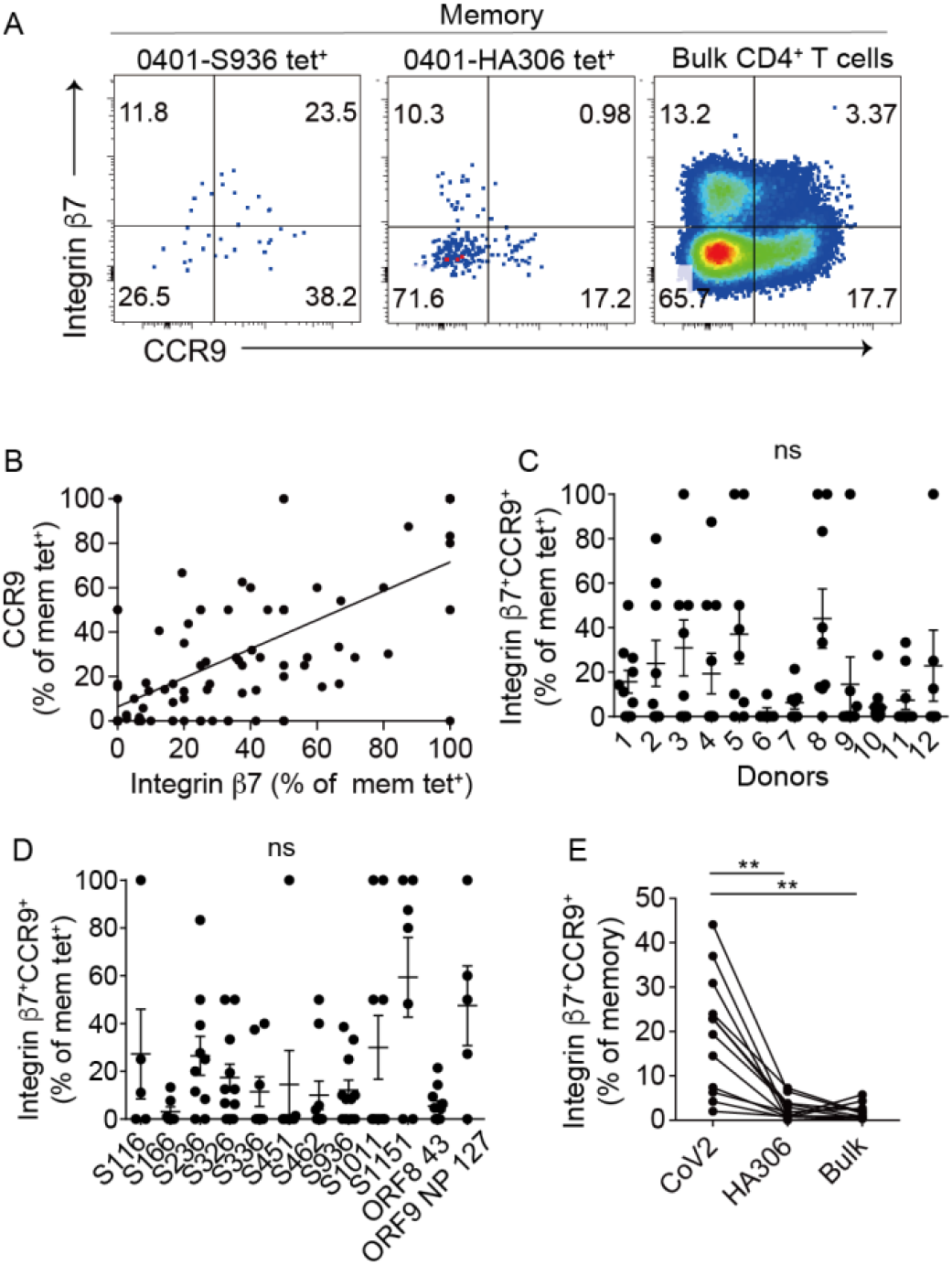
SARS-CoV-2 specific T cells express gut-trafficking receptors. (A) Representative integrin β7 and CCR9 staining on tetramer-labeled and background CD4^+^ T cells. Naïve cells were excluded. (B) The relationship between integrin β7 and CCR9 expression on SARS-CoV2 tetramer^+^ memory cells. Each symbol represents data from a distinct SARS-CoV-2-specific population (n = 102 populations). (C -D) The abundance of integrin β7^+^CCR9^+^ expression as a percentage of SARS-CoV2 tetramer^+^ memory cells by donor (C) or specificity (D). (E) Integrin β7^+^CCR9^+^ frequency as a percentage of memory cells that recognized peptides from SARS-CoV-2 or influenza virus (HA306-318), or as a percentage of memory cells in the tetramer negative fraction. Distinct SARS-CoV-2 tetramer^+^ populations from the same donor are combined and represented as an average. Each symbol represents data from one individual (n = 12). Line connects data from the same donor. For (B) Spearman correlation was computed (0.5984, p< 0.001). Line represents least square regression line. (C) and (D) used Welch’s ANOVA. P-values for pairwise comparisons were computed using Dunnett’s T3 procedure. For (E), repeated measure one-way ANOVA was used and p-values for pairwise comparisons were computed using Tukey’s procedure. Data are shown as Mean ± SEM. ** p < 0.01.

### SARS-CoV-2 specific T cells respond to microbial antigens

While T cells recognize antigens in a highly specific manner, it is known that TCRs can also flexibly dock onto pMHC complexes *(40, 41)*. This feature of the TCR that allows a single TCR to bind multiple distinct pMHCs is referred to as cross-reactivity. Here we investigated whether SARS-CoV-2-specific T cells can cross-react with commensal bacteria-derived antigens. We selected T cells that recognize the spike amino acid sequence 936-952 (S936) to perform in-depth analyses on cross-reactive responses. We focused on this population because S936 specific T-cells showed high integrin β7 and CCR9 expression and were sufficiently abundant to enable single cell cloning. We sorted 96 S936 tetramer^+^ cells into 96-well plates containing irradiated feeder cells and expanded them for 3-4 weeks to generate single cell clones (Fig. 3A). Out of the six clones that grew, three that stained most strongly for tetramers after in vitro expansion were selected for downstream analyses. To identify potential cross-reactivity to microbial peptides, we predicted the DR4 binding register in S936 using NetMHCII 2.3. Protein BLAST performed using S936 core sequence identified six peptides from gut commensal bacteria that shared sequence similarity (Fig. 3B, Table S4). We stimulated S936 clones with peptide-loaded monocyte-derived dendritic cells (DC) and used intracellular TNF-α staining to identify responding T cells. This showed different S936 clones generated distinct responses to commensal microbial peptides in a clone-specific pattern (Fig. 3C, S3A). We also generated tetramers with bacterial peptides and showed that a subset of cross-reactive interactions had sufficient strength to be detected by tetramer staining (Fig. 3D, S3B). The extent of cross-reactive response by tetramer staining correlated with the magnitude of cytokine response by peptide stimulation (Fig. 3E). These data suggest that intestinal microbes have the potential to drive cross-reactive T cell activation. To test how T cells respond to naturally occurring microbial components, S936 clones were incubated with DCs treated with fecal lysates generated from 7 healthy individuals. Among the three clones tested, S936-C3 generated a broad response to different fecal lysates whereas S936-C9 and S936-H6 clones were unresponsive (Fig. 3F and S4A-B). Pre-treating lysate loaded DCs with major histocompatibility complex II (MHC II) blocking antibodies inhibited the production of TNF-α by S936-C3, indicating that cytokine response to fecal lysates was MHC-dependent (Fig. 3G, S4C). These data provide examples of T cell cross recognition between SARS-CoV-2 and other microbial peptides and highlight cross-reactive potential of pre-existing T cells.

**Figure 3:**
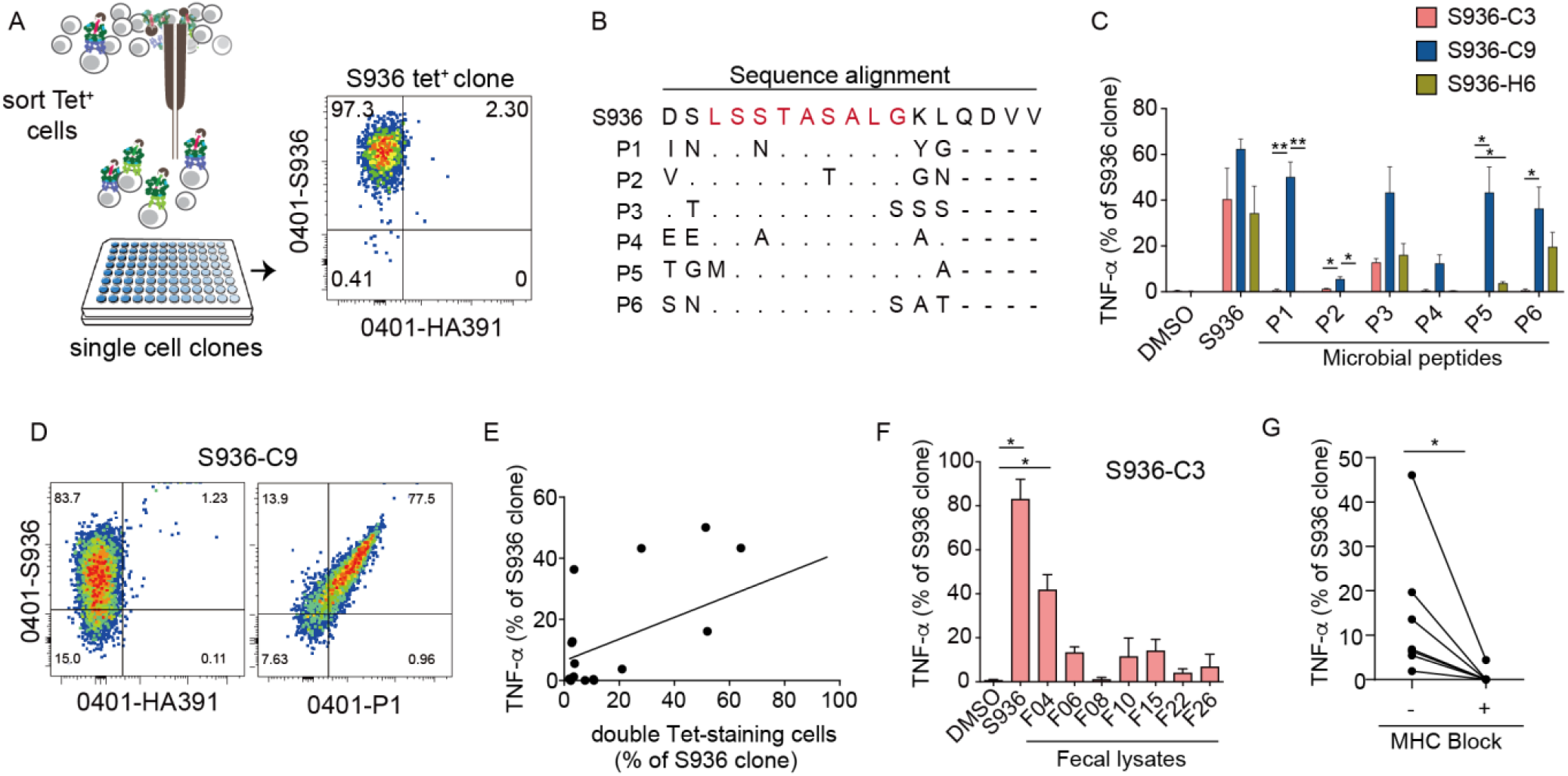
Pre-existing memory to SARS-CoV-2 cross-react with commensal-derived antigens. (A) Schematics illustrating tetramer sorting and in vitro culture to generate single T cell clones. Plot shows representative staining to confirm tetramer-binding of the expanded clone. (B) Sequence alignment of Spike sequence with commensal bacteria-derived peptides. Predicted HLA-DR binding register is colored in red. (C) Plots show T cell response from three S936 clones after an 8-hour stimulation with vehicle, cognate spike peptide, or commensal microbial peptides. Responding T cells are identified by intracellular cytokine staining for TNF-α. (D) Representative plot showing co-staining of S936-C9 clone by tetramers loaded with S936 or P1, a *Bacteroides* TonB-dependent receptor-derived sequence. Tetramers containing a non-cross-reactive influenza peptide was used as a negative control (HA 391-410). (E) The relationship between T cells’ ability to respond to microbial peptides by TNF-α production and to bind the same peptide by tetramer-staining. Each symbol represents measurements from each clone to one microbial peptide. Plot combines data from S936-C3, C9, and H6. (F) The frequency of S936-C3 clone that stained for TNF-α after stimulation with DCs treated with vehicle, cognate peptide, or fecal lysates from 7 healthy adults. (G) The frequency of TNF-α^+^ cells from S936-C3 clone that responded to fecal lysates in the absence or presence of anti-MHC class II blocking antibodies. Each symbol represents treatment with a different fecal lysate. For (C), a two-way ANOVA was used with p-values for pairwise comparisons computed using Tukey’s procedure. For (E) Pearson correlation was computed (0.5589, p = 0.0159). Line represents least square regression line. For (F) Welch’s ANOVA was used with p-values for pairwise comparisons computed using Dunnett’s T3 procedure. For (G), paired t-test was used. Data are shown as Mean ± SEM. * p < 0.05, ** p < 0.01.

### Pre-existing SARS-CoV-2 tetramer^+^ cells are phenotypically heterogeneous

To further investigate the differentiation state and phenotypic diversity of SARS-CoV-2 specific T cells at baseline, we designed a 27 fluorochrome spectral cytometry panel that focused on trafficking receptor expression. We pooled twelve SARS-CoV-2 tetramers on the same fluorochrome to maximize the capture efficiency of SARS-CoV-2-specific T cells from a limited amount of pre-pandemic blood sample. Tetramers loaded with an influenza HA peptide were included for comparison. Tetramer staining, enrichment, and co-staining with cell surface markers were performed using PBMCs from six healthy donors. SARS-CoV-2 tetramer labeled T cells from each donor were identified by manual gating and combined for the Spectre analysis pipeline *(42)*. Five immune clusters were identified using Phenograph and visualized in two-dimensions by Uniform Manifold Approximation and Projection (UMAP) (Fig. 4A) *(43, 44)*. Consistent with our lower dimensional data in previous figures, we observed cells that stained for CD45RO, integrin β7, and CCR9 on the UMAP (Fig. S5). A portion of CD45RO^+^ cells also stained for Tfh-associated marker, CXCR5, and Th1-associated marker, CXCR3, suggesting polarization of some pre-existing memory cells into defined T helper subsets (Fig. S5). Unexpectedly, we also detected SARS-CoV-2 tetramer^+^ cells that express skin homing markers, cutaneous lymphocyte-associated antigen (CLA) and CCR10, which localized to regions that were largely negative for Integrin β7 staining (Fig. S5) *(45-47)*. We then generated a heatmap to examine the composition of Phenograph-defined clusters (Fig. 4B). This showed that cells were primarily partitioned based on trafficking potential and differentiation states into a skin-tropic cluster (CLA^+^CCR10^+^, cluster 2), a gut-tropic cluster (Integrin β7^+^CCR9^+^, cluster 5), and a Treg-like population that displayed high CD25 and low CD127 staining (cluster 4). The two remaining clusters displayed naïve cell features and had heterogeneous levels of trafficking receptor expression (CD45RO^-^CCR7^+^, clusters 1 and 3) (Fig. 4B). All clusters were represented in cells from each donor although the abundance varied across individuals (Fig. 4C).

**Figure 4:**
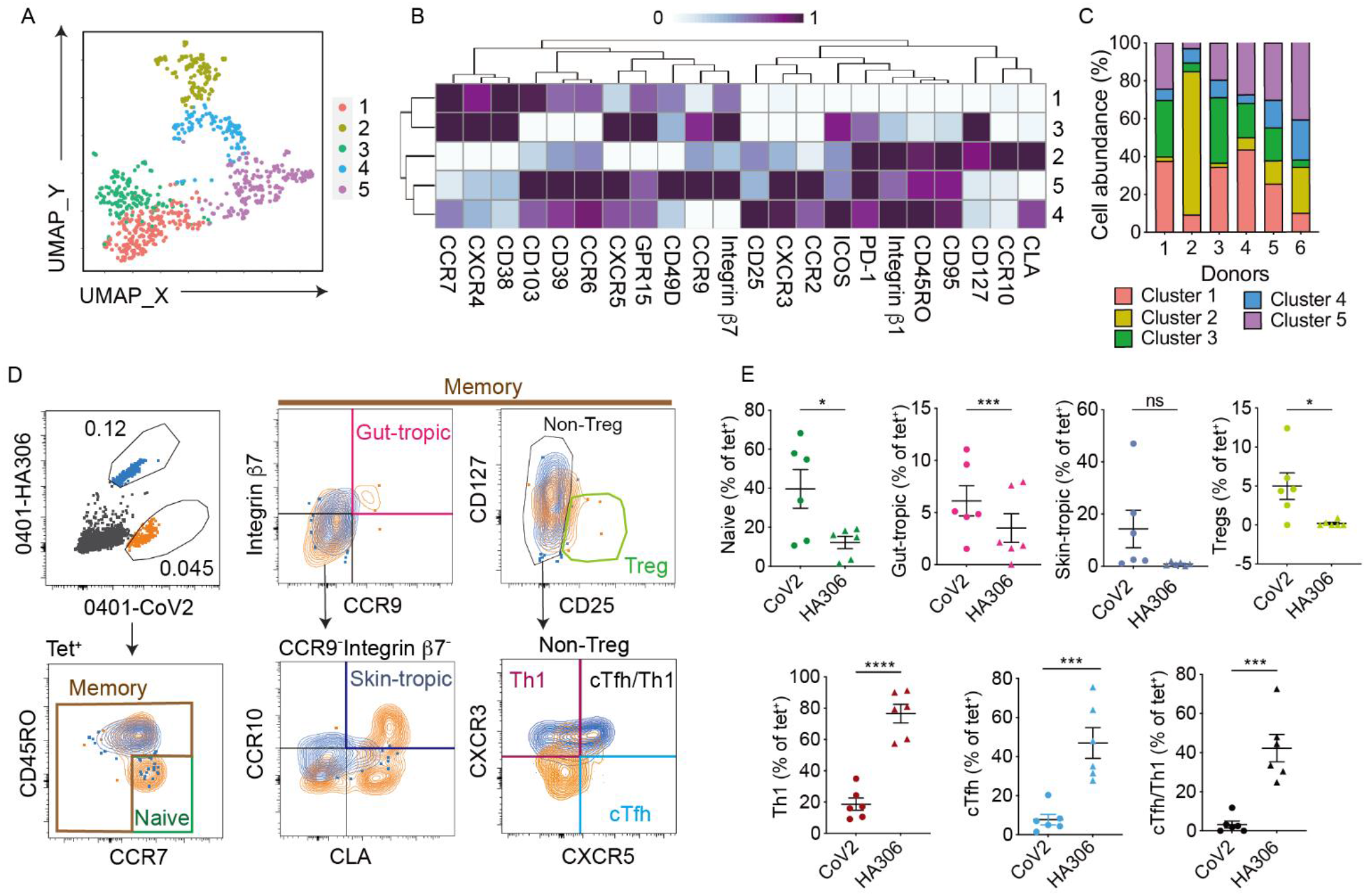
SARS-CoV2 specific T-cells in unexposed donors are phenotypically heterogenous. (A) UMAP displays SARS-CoV-2 tetramer^+^ T cell clusters defined using Phenograph. Data combine 798 cells stained by a pool of 12 SARS-CoV-2 tetramers from 6 healthy individuals. (B) Heatmap shows the median staining signal of the indicated markers for the five SARS-CoV-2-specific clusters shown in A. (C) Bar-graph shows the relative cluster abundance within tetramer^+^ cells from each donor. (D) Representative plots show the gating path to identify tetramer-labeled T cells and each phenotypic subset. (E) Plots summarizes the abundance of indicated subsets within SARS CoV-2 (orange) and HA306 (blue) tetramer-labelled CD4^+^ T cells. Naïve: CD45RO^-^CCR7^+^, gut-tropic: Integrin β7^+^CCR9^+^, skin-tropic: CCR10^+^CLA^+^, Tregs: CD25^high^CD127^low^, Th1: CXCR3^+^, cTfh: CXCR5^+^, cTfh/Th1: CXCR5^+^CXCR3^+^. Each symbol represents data from one donor. For (E), paired t-tests were performed. Data are shown as Mean ± SEM. * p < 0.05, *** p < 0.001 and **** p < 0.0001.

Next, we performed manual gating on tetramer-labeled T cells, focusing on key trafficking, differentiation, and lineage-associated markers to separate out specific subsets (Fig. 4D). Compared to a post-immune HA-specific population, SARS-CoV-2 specific T cells expressed higher frequency of CD45RO^-^CCR7^+^ naïve cells in unexposed individuals (Fig. 4D-E). Within the memory subset, SARS-CoV-2 specific T cells contained more gut-tropic cells compared to HA-specific T cells. Skin-tropic SARS-CoV-2 tetramer^+^ T cells were also detected in a few individuals (Fig. 4D-E). Next, we evaluated cell fate decisions that impact anti-viral immunity. We focused on three key subsets: Tregs, as they can dampen immune responses *(48)*, Tfh cells because they orchestrate B cell response *(49)*, and Th1 cells for direct anti-viral effect, CD8^+^ T cell support, and macrophage activation *(50-52)*. Tetramer^+^ cells were manually gated to identify Tregs by high CD25 and low CD127 expression. Non-Treg cells were further subdivided by CXCR3 and CXCR5 to identify circulating Tfh cells (CXCR5^+^, cTfh), Th1 cells (CXCR3^+^), and cTfh/Th1 cells (CXCR5^+^CXCR3^+^). This showed a distribution of CD4^+^ subsets within SARS-CoV-2-specific T cells that differed from that of a post-immune population. Compared to HA-specific T cells, SARS-CoV-2 tetramer-labeled populations contained a higher frequency of Tregs and fewer Tfh and Th1-skewed T cells (Fig. 4D-E). Collectively, these data revealed a highly heterogeneous pre-existing repertoire to SARS-CoV-2 in unexposed individuals. T cell priming likely originated beyond the gastrointestinal tract to involve other barrier sites such as skin and resulted in a pre-existing memory population that exhibit diverse range of trafficking potential and polarization states.

## Discussion

T cells that recognize SARS-CoV-2 have been found in unexposed individuals *(15, 19-25)*, but the composition of these cells remain poorly defined. Here we directly examine the pre-existing state of 117 SARS-CoV-2 specific T cell populations in healthy adults using pre-pandemic blood samples. The tetramer-based enrichment approach provided the sensitivity necessary to detect and characterize rare antigen-specific T cells directly ex vivo, avoiding potential changes to cellular phenotypes from in vitro cultures. Our analyses of the precursor repertoire to SARS-CoV-2 showed that over 60% of SARS-CoV-2 specific T cells have acquired a memory phenotype prior to known exposure. The naïve fraction of SARS-CoV-2 tetramer^+^ cells was in the lower range of what we had previously detected for YFV-specific T cells in YFV naïve individuals. However, the difference was modest. The presence of pre-existing memory T cells being largely independent of exposures to related pathogens suggests a broader influence that drive precursor differentiation.

Prior studies from Powrie and colleagues found that commensal reactive CD4^+^ T cells primarily expressed a memory phenotype, suggesting that non-infectious stimuli can promote T cell differentiation *(36)*. Based on this and the established close interactions between humans and the microbial environment where the gut microbiota has been shown to critically influence host immunity *(53)*, we hypothesized that the gut microbiota shapes the pre-existing repertoire of SARS-CoV-2-specific T cells. In support of this idea, we identified SARS-CoV-2 specific T cells that expressed gut-homing receptors and cross-reacted with commensal-derived peptides and fecal lysates. While we did not extensively survey the breadth of antigen-recognition, our data provided a well-defined example of cross-reactivity for SARS-CoV-2 precursor T cells. The source of microbial experience is likely to extend beyond intestinal compartment and include exposures that occur at other barrier surfaces. In addition to gut-trafficking, we also identified skin-tropic SARS-CoV-2-specific T cells that expressed CCR10 and CLA. These data suggest that precursor T cells have access to a broad range of tissue environments. T cell experiences with antigens in barrier tissues may provide critical signals that shape precursor repertoire to a novel pathogen.

The composition of the pre-immune repertoire directly impacts post-immune responses. We had previously examined T cell responses in primary immunization with YFV vaccine and identified dynamic changes after vaccination that depended on cells’ pre-existing state *(18)*. Here we performed high dimensional phenotypic analyses on tetramer^+^ cells to further delineate the differentiation states of SARS-CoV-2 specific precursor cells. We identified phenotypic subsets with CD25highCD127low, CXCR5, or CXCR3 expression that suggested regulatory and effector polarization. Finding Tregs within pre-existing populations is consistent with their potential to cross-recognize microbial antigens and the well-established role that gut commensal bacteria plays to drive Treg differentiation *(54)*. Our prior work had also identified Tregs in other virus-specific precursor populations and together they provide evidence for a regulatory component in the baseline response to pathogens *(55)*. We also found CXCR5 and CXCR3-expressing SARS-CoV-2 specific T cells, but they represented a small subset of the precursor repertoire. How pre-existing memory cells influence post-exposure response remain incompletely understood. Emerging studies are showing an association between the abundance of pre-existing T cells and beneficial immune responses *(26-28)*. We speculate that the expression of trafficking receptors could enable accelerated migration of pre-conditioned memory T cells into tissues to enhance local defense in the event of infection. The small populations of pre-existing Tfh and Th1 cells could orchestrate B cells and other cellular arms of immune defenses to further jump start responses to a pathogen. Alternatively, inappropriately regulated cells that infiltrate tissues could contribute to excessive inflammation and immunopathology. Histological analyses of tissues from deceased COVID-19 patients have identified immune cells, including T cells, in the inflamed lung parenchyma *(56)*. Our data highlight a distinct pre-existing memory pool poised to engage infectious challenges. It remains to be determined how different pre-existing populations and baseline polarization states combine to modulate the quality of immune response to pathogens.

In summary, our analyses of SARS-CoV-2 precursor repertoire show that pre-existing memory T cells express diverse phenotypes and display broad tissue tropism. Our data argue for a contribution by non-infectious microbes in the education of the precursor repertoire. Inter-individual differences in precursor abundance and differentiation states could contribute to heterogeneity of human responses to vaccines and infections.

## Material and Methods

### Study Design

The goal of the study was to define the pre-existing state of SARS-CoV-2 specific T cells. Cryopreserved cells were stored from past collection from the Stanford Blood Bank or prior studies at the University of Pennsylvania *(18)*. Subject characteristics are shown in Table S1. Stool samples were baseline samples from a controlled feeding study in healthy adult volunteers (aged 18-60 years) *(57)*. All samples were de-identified and obtained with IRB regulatory approval from the University of Pennsylvania.

### Direct ex vivo T cell analyses and cell sorting

Tetramer staining was carried out as previously described *(16, 18)*. In brief, 30 to 90 million CD3 or CD4 enriched T cells were stained at room temperature for 1 hour using 5 ug of each tetramer in 50ul reaction. Tetramer tagged cells were enriched by adding anti-PE and/or anti-APC magnetic beads and passing the mixture through a magnetized column (Miltenyi). The tetramer-enriched samples were stained with live/dead dyes, exclusion markers (anti-CD19 and anti-CD11b, BioLegend), and other surface markers (Table S5) for 30 minutes at 4°C. Samples were acquired by flow cytometry using LSRII (BD) or sorted on FACS Aria (BD). Frequency calculation was obtained by mixing 1/10^th^ of sample with 200,000 fluorescent beads (Spherotech) for normalization *(16)*. Non-zero populations were included in the analyses performed using FlowJo (BD). Spectral flow cytometric analyses were performed with following modifications: 2ug of tetramers loaded with each of the twelve SARS-CoV2 peptides used in this study were tagged to the same fluorochrome and combined in the staining reaction. Tetramer enriched cells were stained with live/dead dyes, exclusion markers, and a panel of additional surface antibodies (Table S5) for 1h at 4°C followed by fixation with 2% paraformaldehyde. Samples were acquired on spectral flow Cytex AURORA (ARC 1207i).

### Generation and stimulation of T cell clones

Tetramer-stained cells were sorted individually into 96-well plates containing 10^5^ irradiated PBMCs and 10^4^ JY cell line (ThermoFisher) per well in the presence of PHA (1:100, ThermoFisher) and 25ng/ml of IL-7 and IL-15 (PeproTech). IL-2 (50 IU/ml, PeproTech) was added on day 5 and replenished every 3-5 days for 3-4 weeks. To stimulate the T cell clones, peripheral monocyte derived dendritic cells (DC) were generated according to McCurley et al. and matured overnight in LPS (100 ng/ml) with vehicle, 20ug/ml of peptides, or 50ug/ml of fecal lysates *(58)*. Fecal lysates were generated by diluting 100mg of fecal sample in 1ml phosphate buffered saline (PBS) and sonicated 3 times for 1 minute at 4°C with 30 seconds of rest in between. Samples were quickly spun down and protein concentration in the supernatants was quantified by BCA protein assay (Thermo scientific). Co-cultures were performed by adding T cell clones that had rested overnight in fresh media without IL-2 to wells containing DCs. Cells were incubated for 8 hours in the presence of monensin (2uM, Sigma) and Brefeldin A (5ug/mL, Sigma). For MHC blocking experiments, anti-MHC class II antibodies, L243 (0.37ug/ul, BioLegend) and TU39 (0.01ug/ul, BioLegend), were added to T cells before combining the cells with DCs. After stimulation, intracellular cytokine staining with anti-TNF-α (BioLegend) was performed using BD Cytofix/Cytoperm Fixation/Permeabilization Kit according to manufacturer protocol (BD).

### Sequence similarity calculation

All spike and non-spike sequence of SARS-CoV2 and common corona viruses were derived from NCBI database. (SARS-CoV2 spike: YP_009724390.1; SARS-CoV2 ORF8: YP_009724396.1; SARS-CoV2 NP: QSM17284.1; HKU1 spike: ABD96198.1; HKU1 ORF8: AZS52623.1; HKU1 NP: YP_173242.1; 229E spike: AWH62679.1; 229E NP: P15130.2; NL63 spike: APF29063.1; NL63 NP: Q6Q1R8.1; OC43 spike: QEG03803.1; OC43 NP: P33469.1). Pair-wise alignment using Clustal Omega was performed to identify aligned regions. Similarity between the aligned sequences were calculated by Sequence Identity And Similarity (SIAS) using BLOSUM62 matrix. A score of 0 was used if no aligned region was found. Missing sequences were excluded from the analyses.

### High-dimensional phenotypic analyses

Spectral cytometric data were analyzed by manual gating to select SARS-CoV-2 tetramer-labeled cells from each sample. Tetramer^+^ cells were exported and computational analyses were performed using the Spectre package in R *(42)*. A total of 798 tetramer^+^ cells were read into R by flowCore and combined into one single dataset for the following data processing and high-dimensional analyses. Staining intensities were converted using Arcsinh transformation with a cofactor of 2000. Batch alignment was performed by first coarse aligning the batches with quantile conversions of marker intensities calculated with a reference sample included in all the batches and then applying this conversion to all samples via the CytoNorm algorithm. Clustering was performed using Phenograph with nearest neighbors set to 60 (k = 60) *(44)*. Unbiased uniform manifold approximation and projection (UMAP) was used for dimensional reduction and visualization *(43)*.

### Statistical Methods

Data transformation was performed using logarithmic function or inverse hyperbolic sine transformation 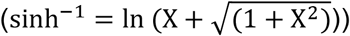 when data contained zero values. Assessment of normality was performed using D’Agostino-Pearson test. Spearman was used if either of the two variables being correlated was non-normal. Otherwise, Pearson was used to measure the degree of association. The best-fitting line was calculated using least squares linear regression. Statistical comparisons were performed using two-tailed Student’s t-test or paired t-test using a p-value of <0.05 as the significance level. Multiple comparisons were performed when the Welch’s one-way ANOVA, repeated measures one-way ANOVA, two-way ANOVA, or mixed effect model was significant. P-values were adjusted for multiple comparisons. Statistical analyses were performed using GraphPad Prism. Lines and bars represent mean and variability is represented by standard error of the mean (SEM). * P < 0.05, ** P < 0.01, *** P < 0.001, **** P < 0.0001.

## Acknowledgments

We thank Mark M. Davis for enabling the collection of cells from healthy blood donors at Stanford. We also thank Annabel Sangree for helpful discussions.

## Author contributions

Conceptualization, L.F.S.; Experimentation, L.B., S.A., Y.P., L.W, C.L, E.S.F.; High-dimensional data analyses, R.X.; Statistics, P.A.G; Supervision, L.F.S., G.D.W; Manuscript preparation, L.F.S., L.B., Y.P., S.A. and R.X.

## Material and Data availability

This study did not generate new unique reagents. All Data are included in the paper.

## Supplementary Material

**Table S1:** Participant information.

| Participant ID | Gender | Sample collection year | Age at the time of collection |
| --- | --- | --- | --- |
| 1 | M | 2017 | 42 |
| 2 | F | 2019 | 52 |
| 3 | M | 2017 | 21 |
| 4 | F | 2017 | 22 |
| 5 | M | 2017 | 25 |
| 6 | M | 2011 | 77 |
| 7 | F | 2011 | 62 |
| 8 | F | 2011 | 61 |
| 9 | M | 2011 | 60 |
| 10 | M | 2011 | 37 |
| 11 | M | 2011 | 70 |
| 12 | M | 2011 | 32 |

**Table S2:**
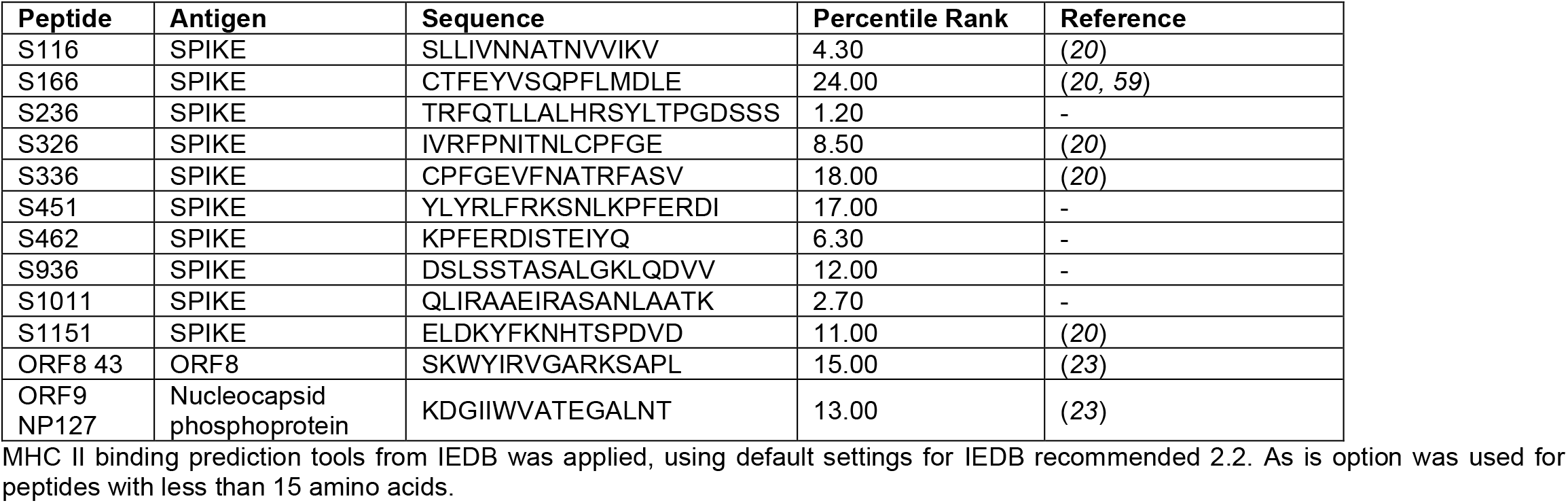
SARS-CoV-2 peptide sequences.

**Table S3:** Sequence alignment between SARS-CoV-2 and common circulating coronaviruses.

|  | S116 | Score |
| --- | --- | --- |
| SARS-CoV-2 | 116 S L L I V N N A T N V V I K V 130 |  |
| HKU1 | 136 T I V . G S V F I . N S Y T I 150 | 40.0 |
| OC43 | 155 G V N K L Q G L L E . S V C Q 169 | 33.3 |

|  | S462 | Score |
| --- | --- | --- |
| SARS-CoV-2 | 462 K P F E R D I S T E I Y Q 474 |  |
| HKU1 | 474 C . C A K P S F A S S C K 486 | 15.4 |
| NL63 | 450 C . . S F S K L N N F Q K 462 | 15.4 |
| OC43 | 473 . . Q P A G V F . N H D V 485 | 30.8 |

|  | S166 | Score |
| --- | --- | --- |
| SARS-CoV-2 | 166 C T F E Y V S Q P F L M D L E 180 |  |
| HKU1 | 183 E S W H F D K S E P . C L F K 197 | 26.7 |
| 229E | 115 D V I R . N I N F E E N L R R 129 | 13.3 |
| NL63 | 175 . Y . N . S C V F S V V N A T 189 | 26.7 |
| OC43 | 201 . L Y K R N F T Y D V N A T Y 215 | 20.0 |

|  | S936 | Score |
| --- | --- | --- |
| SARS-CoV-2 | 936 D S L S S T A S A L G K L Q D V V 952 |  |
| HKU1 | 1021 N G F . A . N . . . A . I . S . . 1037 | 58.8 |
| 229E | 805 . A F T G V N D . I T Q T S Q A L 821 | 29.4 |
| NL63 | 988 A . F . . V N D . I T Q T A E A I 1004 | 41.2 |
| OC43 | 1029 E G F D A . N . . . V . I . A . . 1045 | 58.8 |

|  | S236 | Score |
| --- | --- | --- |
| SARS-CoV-2 | 236 T R F Q T L L A L H R S Y L T P G D S S S 256 |  |
| HKU1 | 233 S H Y Y V . P L T C N A I S S N T . N E T 253 | 33.3 |
| 229E | 179 . S A F V G A L P K T V R E F V I S R T G 199 | 14.3 |
| NL63 | 249 S . L Y Q P . R . T C L W P V . . L K . . 269 | 42.9 |
| OC43 | 251 S H Y Y V M P L T C I . R R N I . F T L E 271 | 28.6 |

|  | S1011 | Score |
| --- | --- | --- |
| SARS-CoV-2 | 1011 Q L I R A A E I R A S A N L A A T K 1028 |  |
| HKU1 | 1096 . . S D I S L V K F G . A . . M E . 1113 | 44.4 |
| 229E | 894 T . T K Y T . V . . . R Q . . Q Q . 911 | 61.1 |
| NL63 | 1077 V . N K Y T . V . S . R R . . Q Q . 1094 | 50.0 |
| OC43 | 1104 . . S D S T L V K F . . A Q . M E . 1121 | 44.4 |

|  | S326 | Score |
| --- | --- | --- |
| SARS-CoV-2 | 326 I V R F P N I T N L C P F G E - 340 |  |
| HKU1 | 317 V H . R I P D L P D . D I D K . 331 | 20.0 |
| 229E | 289 . . S L . V Y H K - H T . I V L 303 | 20.0 |
| OC43 | 333 V Y . R K P D L P N . N I E A . 347 | 20.0 |

|  | S1151 | Score |
| --- | --- | --- |
| SARS-CoV-2 | 1151 E L D K Y F K N H T S P D V D 1165 |  |
| HKU1 | 1237 . . S H W . . . Q . . I A P N 1251 | 60.0 |
| 229E | 1037 T . Q E L S Y K L P N Y T . P 1051 | 13.3 |
| NL63 | 1227 N . P . . V . P N F D L T P F 1241 | 26.7 |
| OC43 | 1245 . . . Q W . . . Q . . V A P . 1259 | 66.7 |

|  | S336 | Score |
| --- | --- | --- |
| SARS-CoV-2 | 336 C P F G E V F N A T R F A S V 350 |  |
| HKU1 | 327 . D I D K W L . N F N V P . P 341 | 20.0 |
| OC43 | 343 . N I E A W L . D K S V P . P 357 | 20.0 |

|  | ORF8 43 | Score |
| --- | --- | --- |
| SARS-CoV-2 | 43 S K W Y I R V G A R K S A P L 57 |  |
| HKU1 | 76 K E Y P L L T . Y P L L K Q K 90 | 20.0 |

|  | S451 | Score |
| --- | --- | --- |
| SARS-CoV-2 | 451 Y L Y R L F R K S N L K P F E R D I 468 |  |
| HKU1 | 448 . G F N N . N L . S H S V V Y S R Y 465 | 22.2 |
| NL63 | 539 S S W H I Y L . . G T C . . S F S K 556 | 44.4 |
| OC43 | 462 K R F G F I E D . V F . . Q P A G V 479 | 27.8 |

|  | ORF9 NP 127 | Score |
| --- | --- | --- |
| SARS-CoV-2 | 127 K D G I I W V A T E G A L N T 141 |  |
| HKU1 | 141 L E . V F . . . N H Q . D T S 155 | 53.3 |
| 229E | 97 V E . V V . . . V D . . K T E 111 | 66.7 |
| NL63 | 95 S . . V V . . . K . . . K T V 109 | 66.7 |
| OC43 | 142 I . . V Y . . . S N Q . D V N 156 | 53.3 |

**Table S4:**
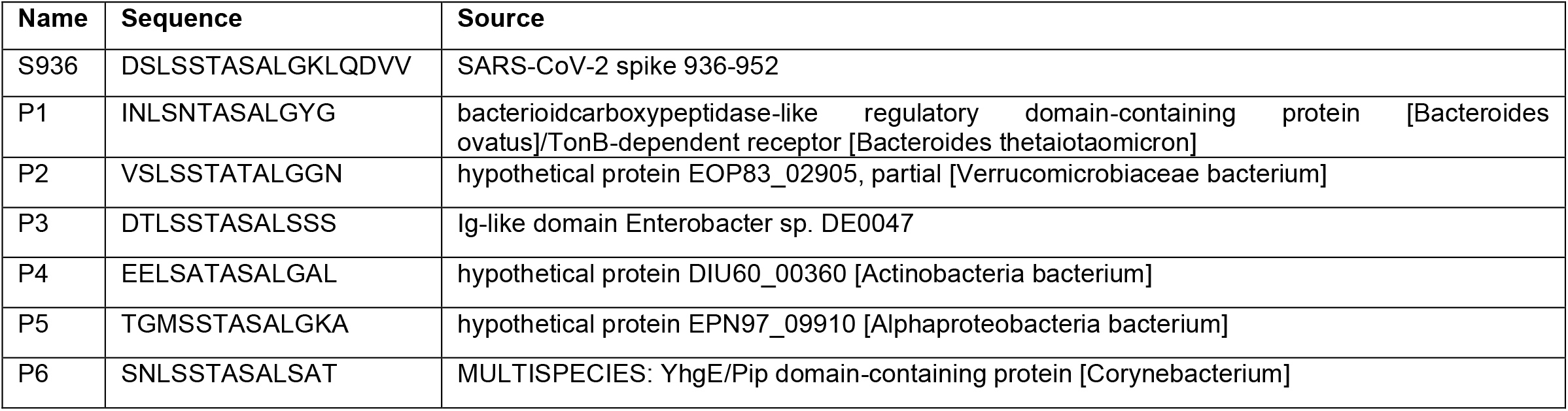
Cross-reactive bacterial peptide sequences.

**Table S5:** Surface antibody staining panels.

| Specificity | Fluorochrome | Clone ID | Flow cytometry |
| --- | --- | --- | --- |
| CD3 | BV785 | UCHT1 | conventional flow |
| CD45RO | BV605 | UCHL1 | conventional flow |
| ICOS | Alexa Fluor 700 | C398.4A | conventional flow |
| CD3 | BUV395 | UCHT1 | Spectral |
| CD45RO | BV570 | UCHL1 | Spectral |
| CD95 | BV711 | DX2 | Spectral |
| CD49D | BV750 | 9F10 | Spectral |
| CD103 | BV785 | Ber-ACT8 | Spectral |
| Integrin b1 | Superbright 600 | TS2/16 | Spectral |
| CCR2 | BUV737 | LS132 | Spectral |
| CCR6 | PerCP-eFluor 710 | R6H1 | Spectral |
| CCR10 | PE | 563656 | Spectral |
| CLA | Pacific Blue | HECA-452 | Spectral |
| GPR15 | BV421 | SA302A10 | Spectral |
| CXCR3 | Alexa Fluor 647 | G025H7 | Spectral |
| CXCR4 | BUV563 | 12G5 | Spectral |
| CXCR5 | BB515 | RF852 | Spectral |
| PD-1 | Alexa Fluor 700 | EH12.2H7 | Spectral |
| CD39 | BV480 | TU66 | Spectral |
| CD127 | Alexa Fluor 532 | eBioRDR5 | Spectral |
| CD25 | APC-Fire 810 | M-A251 | Spectral |
| CD38 | PE-Cy5 | HIT2 | Spectral |
| ICOS | BUV496 | DX29 | Spectral |
| CD4 | BV510 | SK3 | Spectral / conventional flow |
| CCR7 | BV650 | G043H7 | Spectral / conventional flow |
| Integrin b7 | PerCPcy5.5 | FIB504 | Spectral / conventional flow |
| CCR9 | APC | L053E8 | Spectral / conventional flow |
| CD11b | APC-Cy7 | ICRF44 | Spectral / conventional flow |
| CD19 | APC-Cy7 | HIB19 | Spectral / conventional flow |

**Figure S1:**
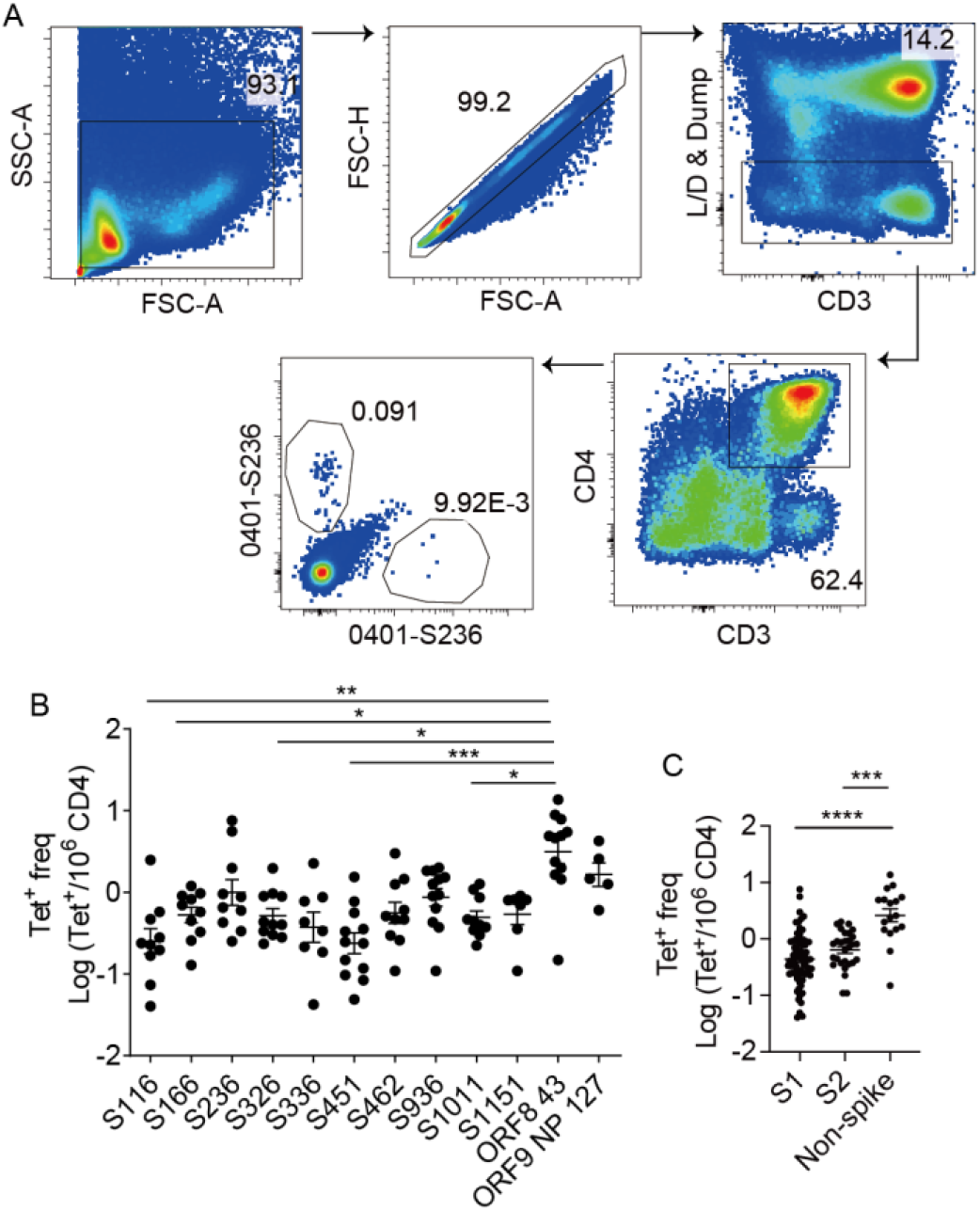
SARS-CoV-2 specific T cell analyses by specificity. (A) Representative plots show the gating strategy for identifying tetramer^+^ cells shown in Fig. 1A. (B) Plot shows the frequency of T cells from different individuals that recognized the same pMHC complex. Each symbol represents data from one donor. (C) Plot summarizes the frequencies of T cells that recognized S1 subunit, S2 subunit, or non-spike regions of SARS-CoV-2. For (B) and (C), Welch’ ANOVA was used with p-values for pairwise comparisons computed using Dunnett’s T3 procedure. Data are shown as Mean ± SEM. * p < 0.05, ** p < 0.01, *** p < 0.001 and **** p < 0.0001.

**Figure S2:**
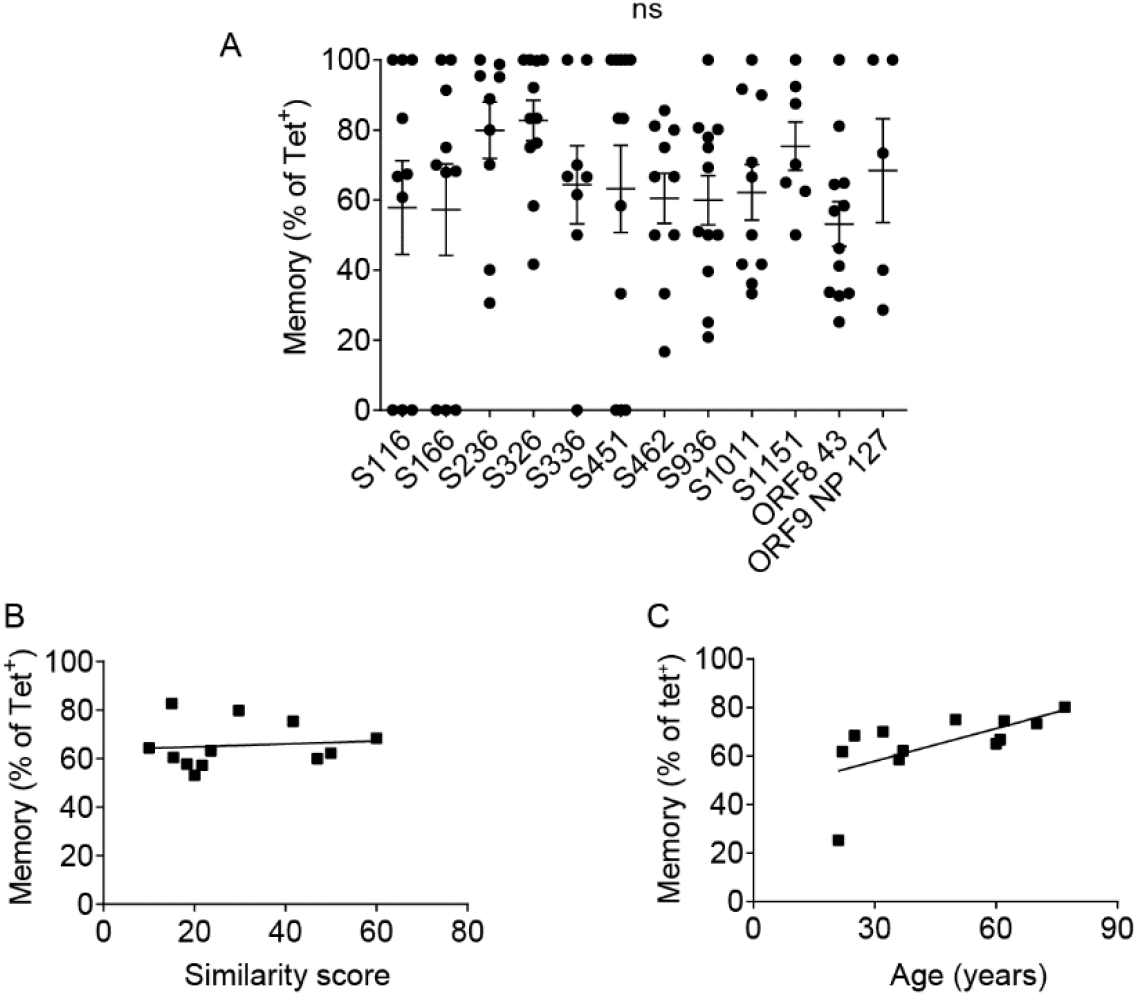
Memory phenotype distribution by specificity, epitope similarity, and donor age. (A) Plot shows the memory phenotype of T cells from different individuals that recognized the same pMHC complex. Each symbol represents data from one donor. (B) Relationship between the proportion of memory cells within a tetramer^+^ population and conservation of sequences with common circulating strains as indicated by the similarity score. Each symbol represents one T cell specificity and combines data from different donors. (C) Relationship between the abundance of memory precursors and donor age. Each symbol represents data from one individual. Distinct SARS-CoV-2 tetramer^+^ populations from the same donor are combined and represented as an average. For (A) Welch’s ANOVA was used with p-values for pairwise comparisons evaluated using Dunnett’s T3 procedure. For (B), Pearson correlation was computed (0.1025, p = 0.7512) For (C), Spearman correlation was computed (0.7063, p = 0.0129). The line represents least square regression line. Data are shown as Mean ± SEM.

**Figure S3:**
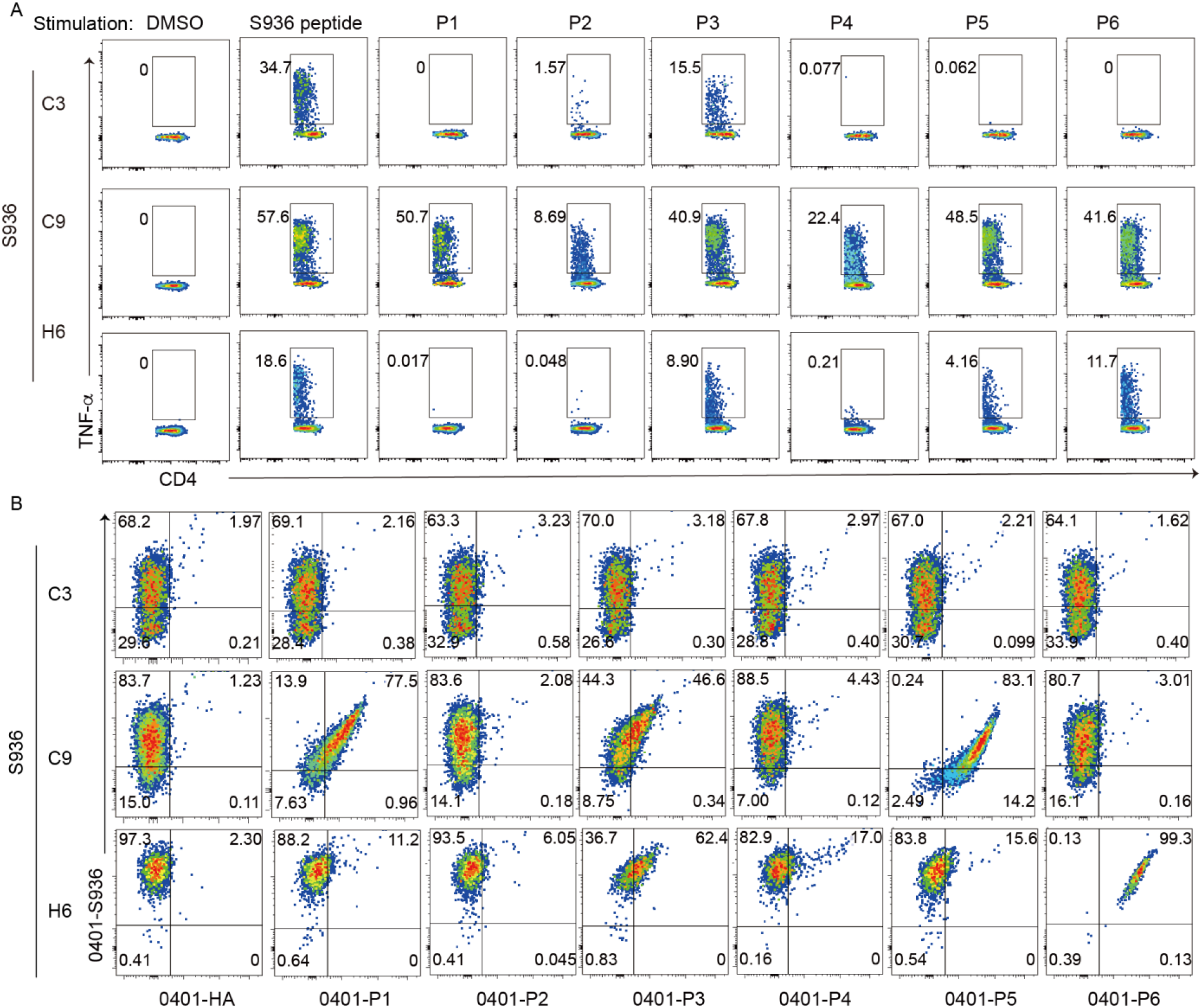
Peptide stimulation and tetramer staining of S936 clones. (A) Plots show representative T cell response generated by each S936 clone after an 8-hour stimulation in the indicated conditions. Responding T cells are identified by intracellular cytokine staining for TNF-α. (B) Plots show tetramer staining of S936 clones with the cognate spike peptide-loaded tetramer (0401-S936) and a second non-spike tetramer. Non-spike tetramers contain a microbial peptide (P1 - P6) or a control influenza peptide (HA 391-410).

**Figure S4:**
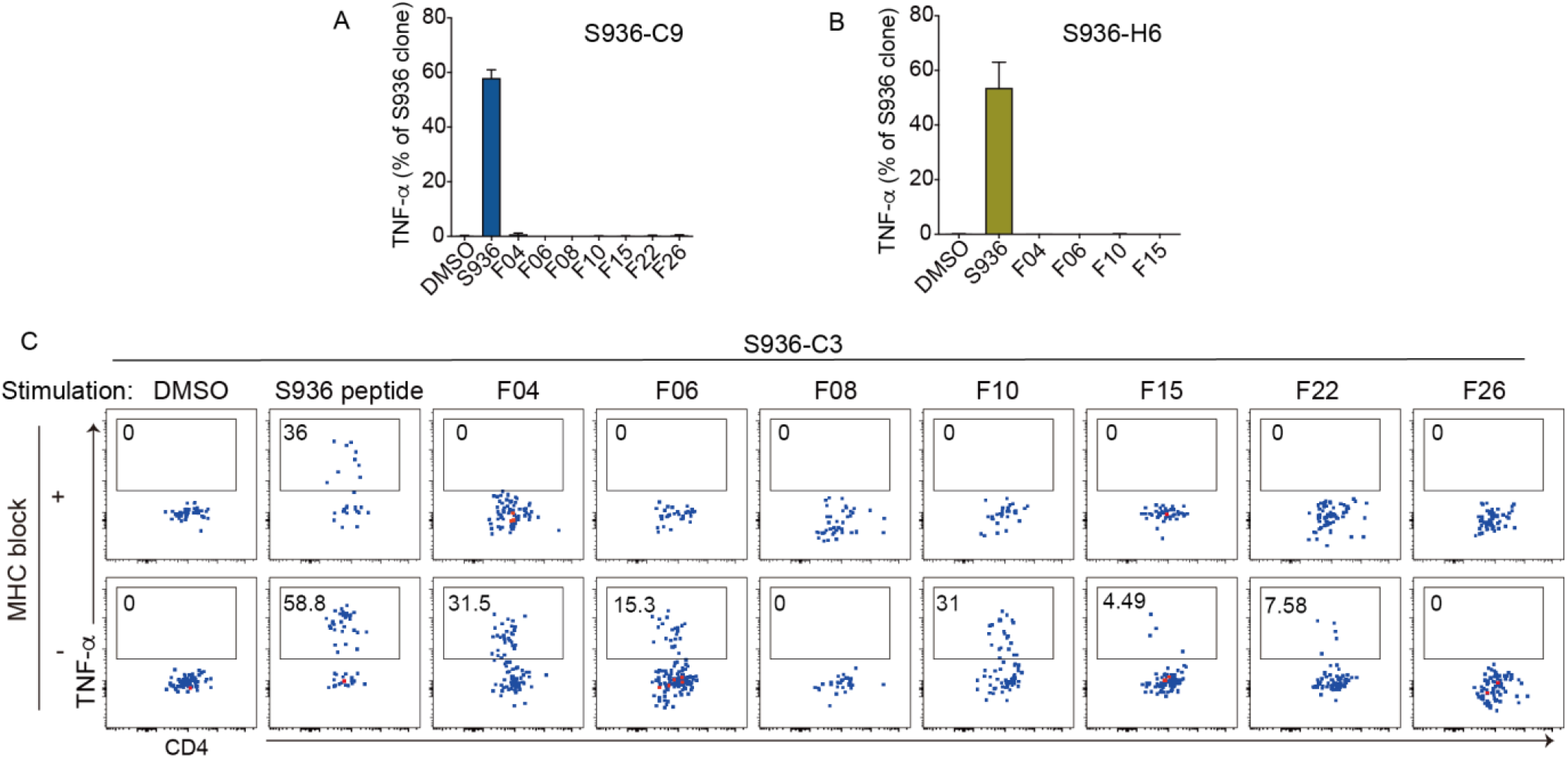
T cell response to fecal lysates. (A-B) The frequency of S936-C9 clone (A) or S936-H6 clone (B) that stained for TNF-α after stimulation with DCs treated with vehicle, cognate peptide, or fecal lysates from healthy adults. (C) Representative plots show TNF-α expression by S936-C3 clone in the indicated conditions, with or without MHC blocking antibodies. Data are shown as Mean ± SEM.

**Figure S5:**
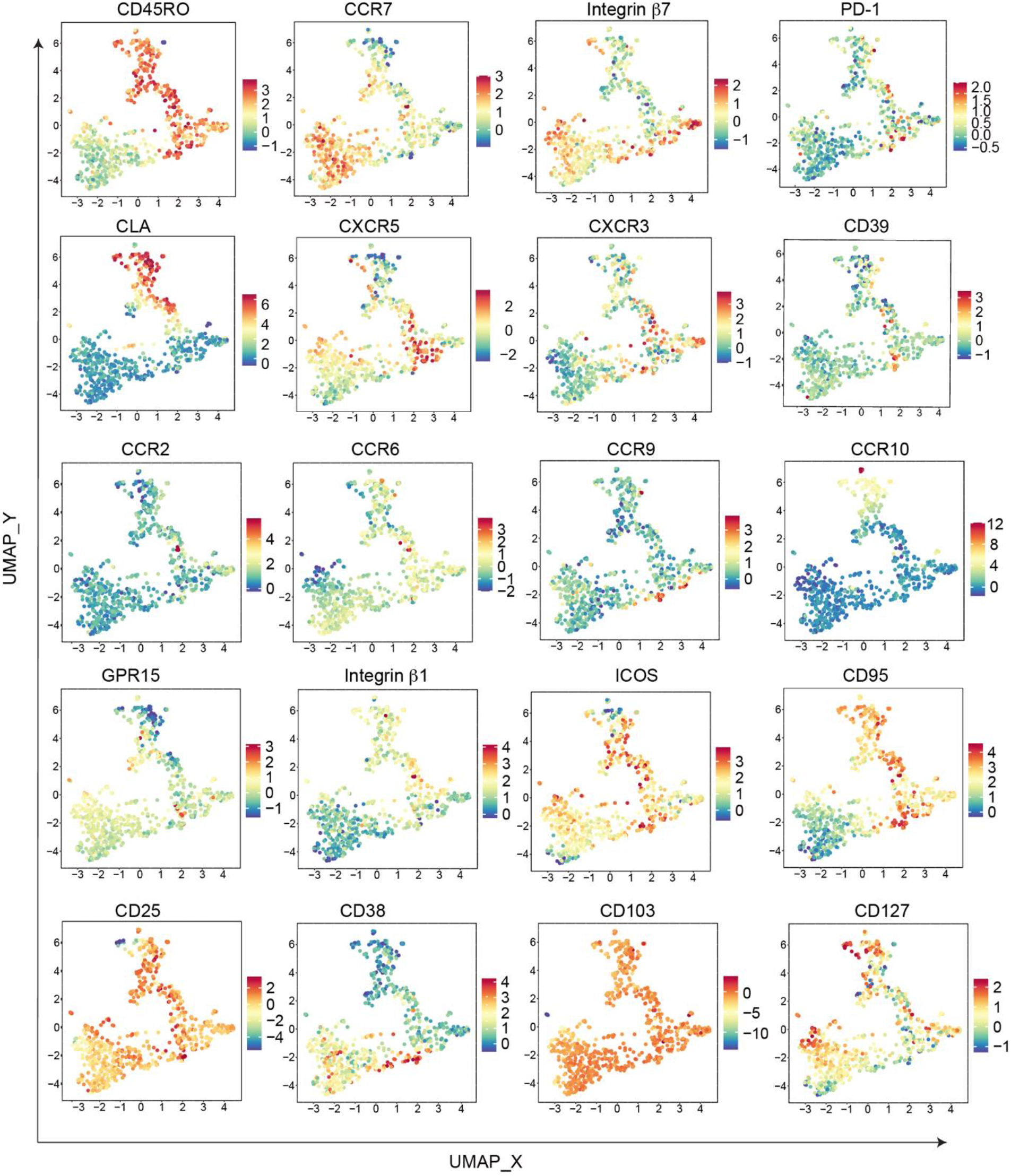
High-dimensional phenotypic analyses of pre-immune SARS-CoV-2 specific T cells. UMAPs display individual markers as indicated. Markers used to select input cells were excluded. Plots combine SARS-CoV-2 tetramer^+^ cells from six individuals.

